# Optogenetic rejuvenation of mitochondrial membrane potential extends *C. elegans* lifespan

**DOI:** 10.1101/2022.05.11.491574

**Authors:** Brandon J. Berry, Anežka Vodičková, Annika Müller-Eigner, Chen Meng, Christina Ludwig, Matt Kaeberlein, Shahaf Peleg, Andrew P. Wojtovich

## Abstract

Despite longstanding scientific interest in the centrality of mitochondria to biological aging, directly controlling mitochondrial function to test causality has eluded researchers. We show that specifically boosting mitochondrial membrane potential through a light-activated proton pump reversed age-associated phenotypes and extended *C. elegans* lifespan. We show that harnessing the energy of light to experimentally increase mitochondrial membrane potential during adulthood alone is sufficient to slow the rate of aging.

## Main

The causal role of mitochondrial dysfunction and metabolic decline are central questions of aging research [1, 2]. The voltage potential across the inner membrane of mitochondria (membrane potential: Δψ_m_) decreases with age in many model systems [3–6]. Δψ_m_ is a fundamental driver of diverse mitochondrial functions, including ATP production, immune signaling, and genetic and epigenetic regulation [7]. Therefore, decreased Δψ_m_ is an attractive explanation for the complex dysfunctions of aging. However, it is unclear whether decreased Δψ_m_ is a cause or a consequence of cellular aging.

To test these questions in a metazoan, we used optogenetics to harness the energy of light using a mitochondria-targeted light-activated proton pump to increase Δψ_m_. Using a mitochondrial targeting sequence, we previously expressed a rhodopsin-related pump that is specific for protons [8] in the inner membrane of mitochondria [9]. We called this tool “mitochondria-ON” or mtON (Figure 1A) and previously fully characterized its optogenetic function [9]. mtON isolates Δψ_m_ as a single experimental variable in vivo, and requires both light activation and a cofactor, all-trans retinal (ATR), for proton pumping activity [9]. *C. elegans* do not produce ATR endogenously, allowing for control conditions of light exposure alone (which can be damaging [10]), mtON protein expression alone, and ATR supplementation alone (which does not affect lifespan or physiology, Supplementary Table 1 and [11]). Only animals supplemented with ATR and illuminated will have mtON activity and increased Δψ_m_ (Supplementary Figure 1) [9].

**Figure 1.**
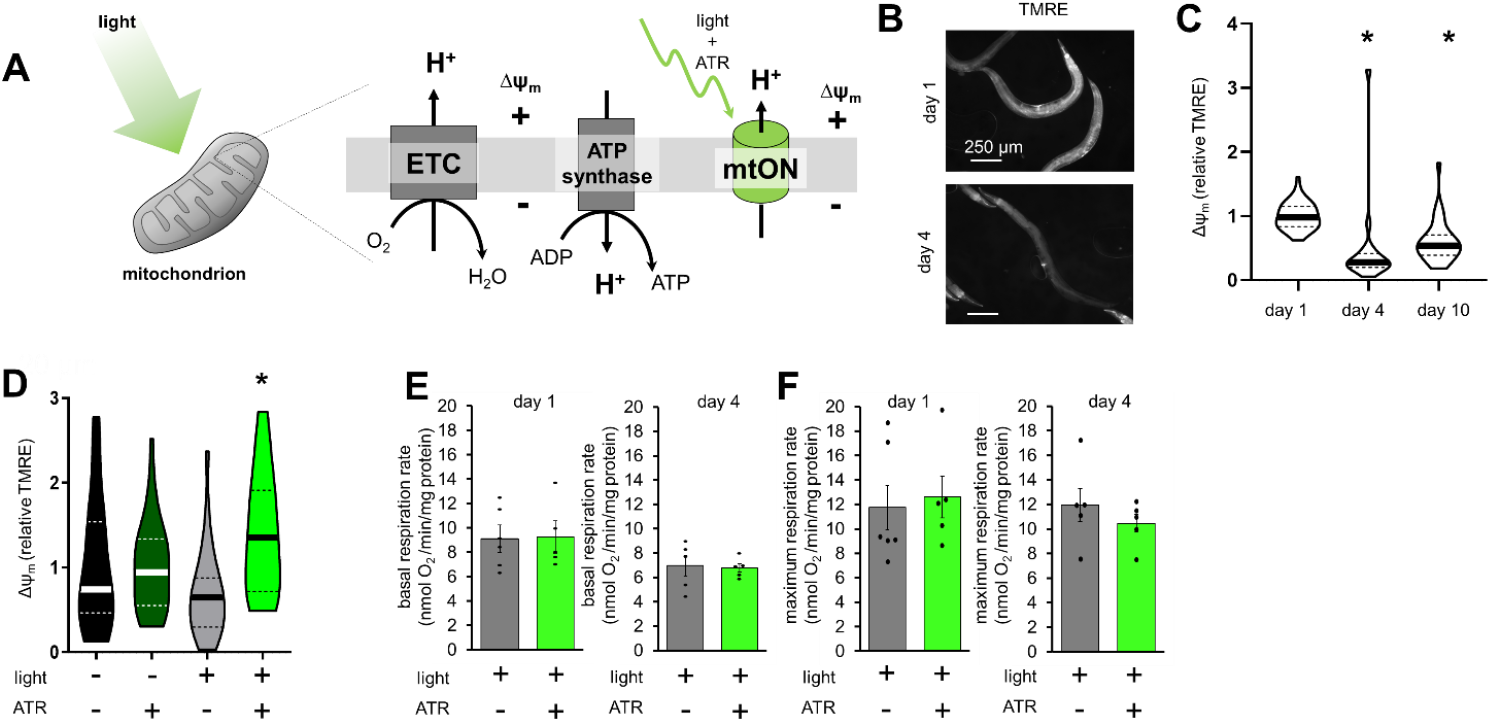
Mitochondria-ON (mtON) increased Δψ_m_ in vivo. A) The mitochondrial inner membrane (IM) contains the electron transport chain (ETC) that pumps protons to generate mitochondrial membrane potential (Δψ_m_). mtON is a light-activated proton pump that contains the cofactor all-trans retinal (ATR), and in response to light, pumps protons across the IM to generate Δψ_m_. B) Adult worms stained with the Δψ_m_ indicator, TMRE. Scale bar is 250 μm. C) Quantification of panel B, relative TMRE fluorescence. One-way ANOVA with Tukey ‘s multiple comparisons test, *p < 0.05, n = 44, 18, 30. Data are medians + quartiles (dotted lines). D) TMRE quantification (representative images in Supplementary Figure 2C). One-way ANOVA with Tukey’s test, *p < 0.05, n = 33, 34, 26, 35 animals for each bar from left to right. Data are medians + quartiles. E) Basal Oxygen consumption of day 1 (left) and day 4 (right) animals, n = 5-6 populations each condition. Data are means + SEM, dots are individual populations. F) Maximal oxygen consumption (induced by FCCP) of the same populations in panel H. Data are means + SEM, dots are individual populations.

We found that Δψ_m_ naturally declines with age in *C. elegans* (Figure 1B&C), as expected [2, 3, 5]. mtON activation reversed that decline (Figure 1D & Supplementary Figure 2) in two different genetic backgrounds (Supplementary Figure 3). mtON activation did not impact mitochondrial mass observed by mitochondrial staining and quantitative proteomics (Supplementary Figure 2A-G & Supplementary Figure 4). Basal respiration rates were similar across conditions (Figure 1E&F) as expected, given mtON’s specificity for Δψ_m_ alone [9].

We activated mtON throughout the lifespan beginning in adulthood and found a reproducible increase in lifespan compared to controls (Figure 2A). Lifespan extension was replicated independently across three different strains, different light intensities, and in different laboratories (Supplementary Table 1), and was sensitive to the mitochondrial uncoupler, FCCP, (Figure 2B) which dissipates Δψ_m_. FCCP had no effect on lifespan on its own (Supplementary table 1). These data indicate that increasing Δψ_m_ causes increased lifespan.

**Figure 2.**
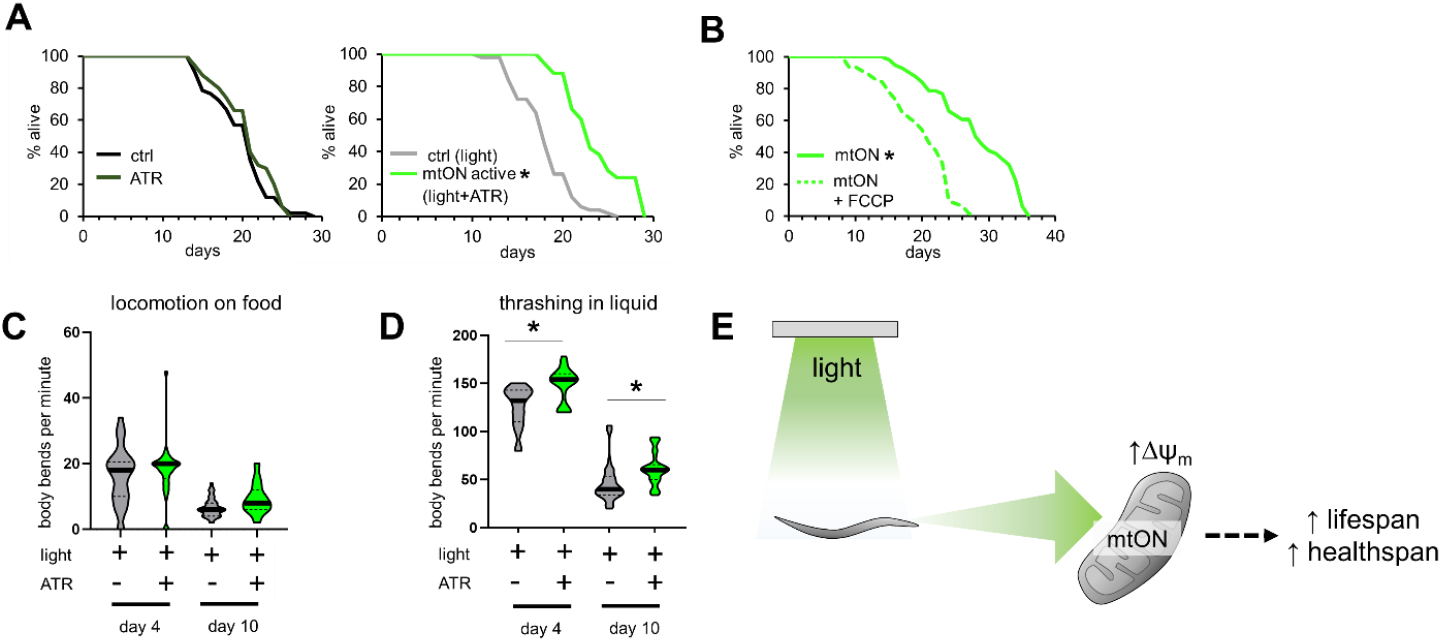
mtON extended lifespan and healthspan. A) Survival curves of mtON-expressing animals. Light treatment began at day 1 of adulthood. Left: ATR supplementation without light had no effect on lifespan by log-rank (Mantel-Cox) test, p > 0.05. Right: mtON activation significantly extended lifespan compared to the light control by log-rank (Mantel-Cox) test, *p < 0.0001. Detailed statistical information is presented in Supplementary Table 1. C) mtON activation (+ATR +light) did not affect locomotion with age on solid media. One-way ANOVA with Tukey ‘s test n = 30 animals for day 4 conditions and 40 animals for day 10. Data are medians + quartiles (dotted lines). D) mtON activation improved thrashing rate in liquid with age. One-way ANOVA with Tukey’s test *p < 0.05. n = 32, 32, 40, 40 animals for each violin from left to right. Data are medians + quartiles. All statistical comparisons are presented in Supplementary Table 2. E) Model showing effects of mtON activation in vivo. The dotted arrow represents the molecular mechanisms to be investigated that link Δψ_m_ to aged physiology.

In *C. elegans*, mild inhibition of mitochondrial function during development (but not during adulthood) extends lifespan [12, 13]; conversely, here we show that attenuating the age-associated decrease in Δψ_m_ in adult animals can extend lifespan. Accordingly, targets of the mitochondrial unfolded protein response did not change after mtON activation (Supplementary Figure 4B). These differences may reflect a lifespan-extending hormetic response from mitochondrial perturbation during development [14–16] versus beneficial effects seen here from directly sustaining mitochondrial function during adulthood [2].

Organisms including humans and *C. elegans* have trouble moving as they age due to physiologic decline [17–19]. This functional decline was mitigated by mtON activation in worms thrashing in liquid, but not on solid media (Figure 2C&D and Supplementary Figure 5). These results show that age-associated physiologic dysfunction can be improved by directly reversing the loss of Δψ_m_ that occurs with age. How improving Δψ_m_ may influence redox metabolites, including NAD^+^/NADH, which are known to impact biological aging, should be further assessed. To begin to probe a potential signaling pathway, we tested the effect of mtON in long-lived worms with constitutively active AMPK signaling [20]. mtON further increased lifespan in this model (Supplementary Figure 6 and Supplementary Table 1), indicating that increased Δψ_m_ can additionally contribute to longevity in parallel of a canonical signaling pathway.

In summary, this study used a novel technology that harnesses the energy of light to generate Δψ_m_ to test the hypothesis that Δψ_m_ causally determines longevity in *C. elegans* (Figure 2E). Prior studies reported that inhibition of mitochondrial function during development can increase lifespan, and our results extend the role of mitochondrial function in aging to adult intervention. Preserved Δψ_m_ during adulthood is sufficient to slow normative aging and improve at least some functional measures of health. This work provides important context for understanding the role of mitochondrial function during aging and suggests the potential of novel approaches to delay aging by using optogenetic technology as an energy replacement source.

## Acknowledgments

BJB is supported by the Biological Mechanisms for Healthy Aging Training Grant NIH/NIA T32 AG066574 and by NIH/NIA grant P30AG013280 to MK. APW is supported by NIH grants (R01 NS092558, R01 NS115906). SP is supported by the DFG grant (458246576) by two Longevity Impetus grants from Norn Group. We also acknowledge the W. M. Keck Microscopy Center and the Keck Center Manager, Dr. Nathaniel Peters for confocal microscopy access and training (NIH S10 OD016240).

## Competing Interests

B.J.B, S.P. and A.P.W. are listed as inventors on a patent application based on some of the work described here.

## Author Contributions

BJB, MK, SP, & APW designed the research. BJB performed the lifespans, imaging and analysis, healthspan experiments, and the data analysis. AV carried out lifespans and respiration experiments. AME assisted with experiments. CM and CL carried out the mass spectrometry. BJB wrote the manuscript with input from MK, SP, & APW.

---

**Supplementary Figure 1.**
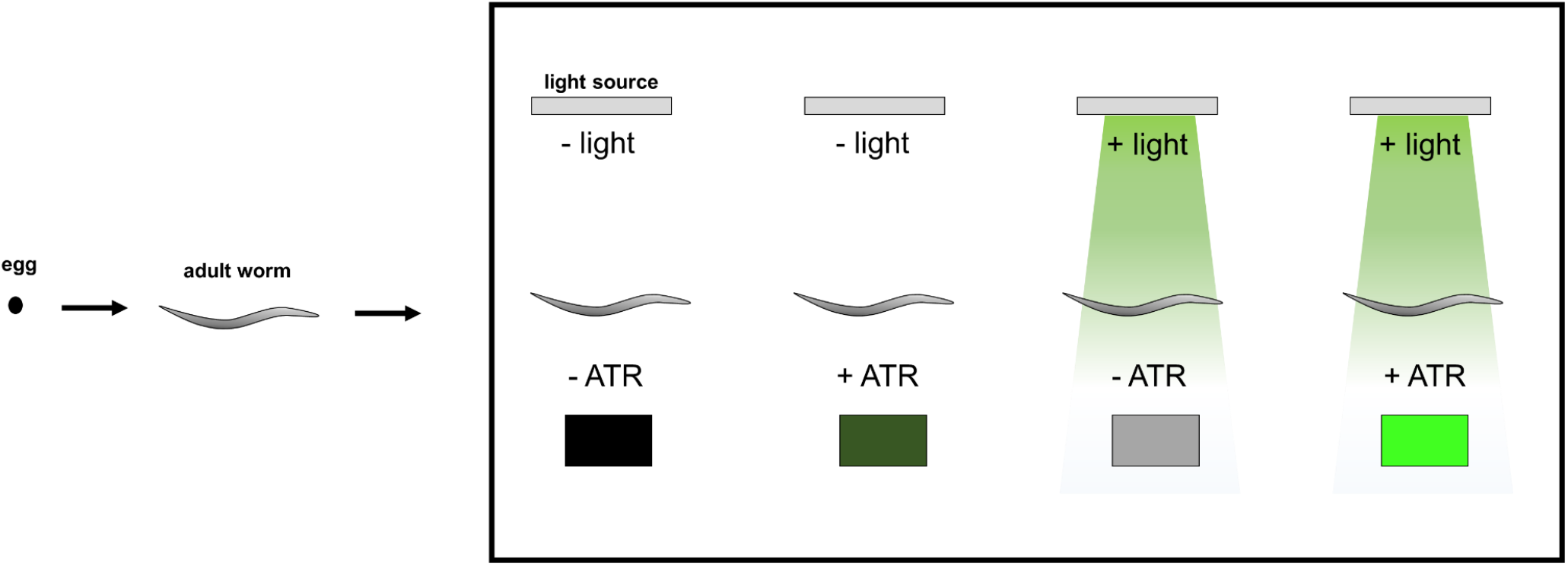
Experimental Schematic. During development, all populations were without light. Starting at day 1 of adulthood, populations were exposed to the indicated conditions until experimental measurements or until death. Light-green color indicates the activated mtON condition (+ light + ATR).

**Supplementary Figure 2.**
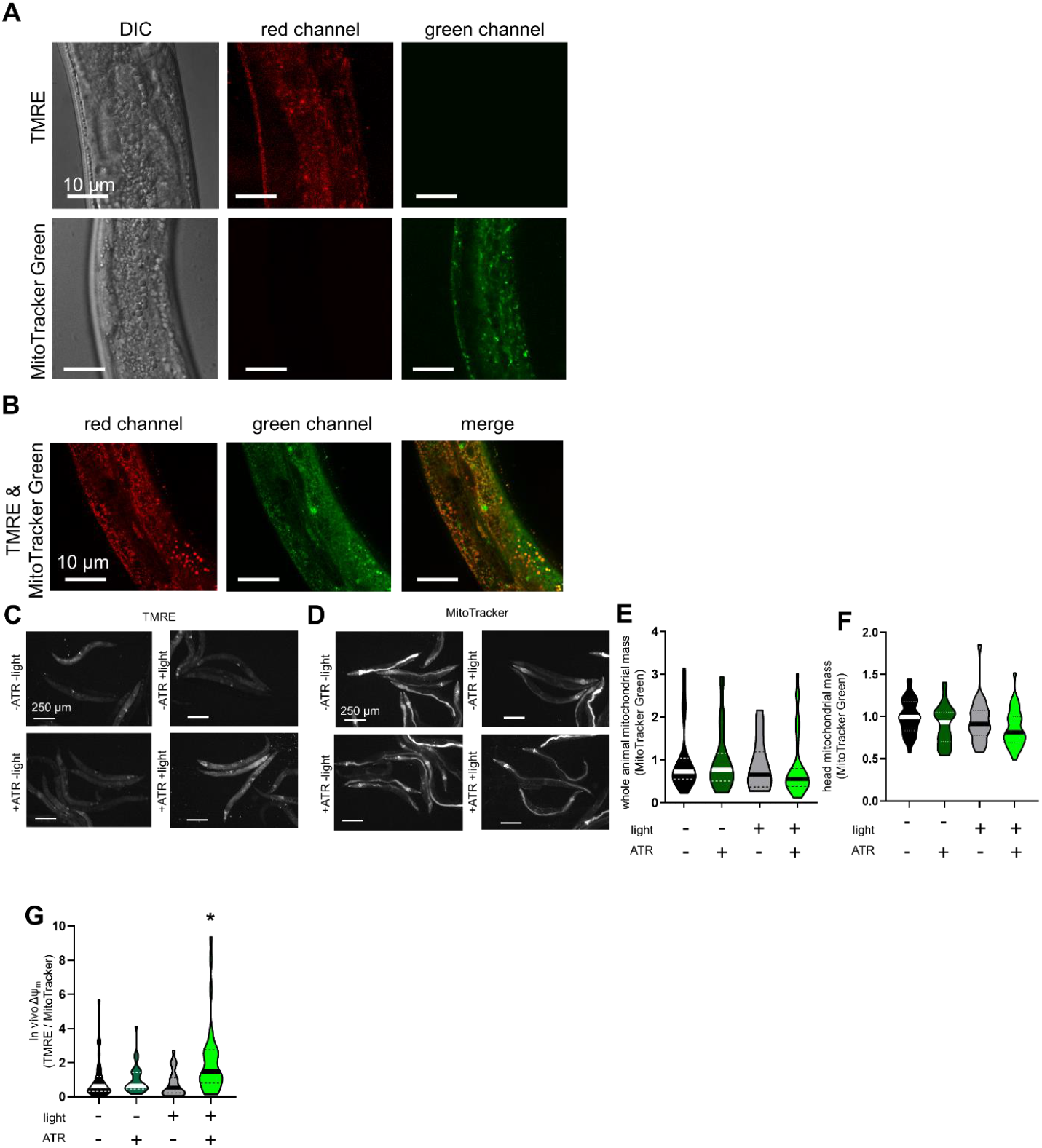
Mitochondrial membrane potential and mass measurements. A) Confocal images of animals stained separately with TMRE or MitoTracker Green, demonstrating mitochondrial patterns in intestine, and showing no channel bleed-through and no fluorescence of extraneous dye in the intestinal lumen. Scale bar is 10 μm. B) Confocal images of animals co-stained with TMRE and MitoTracker Green demonstrating mitochondrial colocalization. Scale bar is 10 μm. *C)* Day 4 adult worms overexpressing mtON stained with TMRE. +ATR +light is the active mtON condition. D) Day 4 adult worms expressing mtON stained with MitoTracker Green. E) Quantification of MitoTracker fluorescence in whole animals (represented in panel D). One-way ANOVA, no significant differences detected between groups. n = 33, 34, 26, 35 animals for each bar from left to right. Data are medians + quartiles. F) Quantification of relative MitoTracker fluorescence in the head regions alone. One-way ANOVA, no significant differences detected between groups. n = 33, 34, 26, 35 animals for each bar from left to right. G) Ratio of TMRE to MitoTracker Green whole-body fluorescence (represented in panels C and D) demonstrating increased mitochondrial membrane potential relative to mitochondrial mass in response to mtON activation. One-way ANOVA with Tukey’s test, *p < 0.05, n = 33, 34, 26, 35 animals for each bar from left to right. Data are medians + quartiles.

**Supplementary Figure 3.**
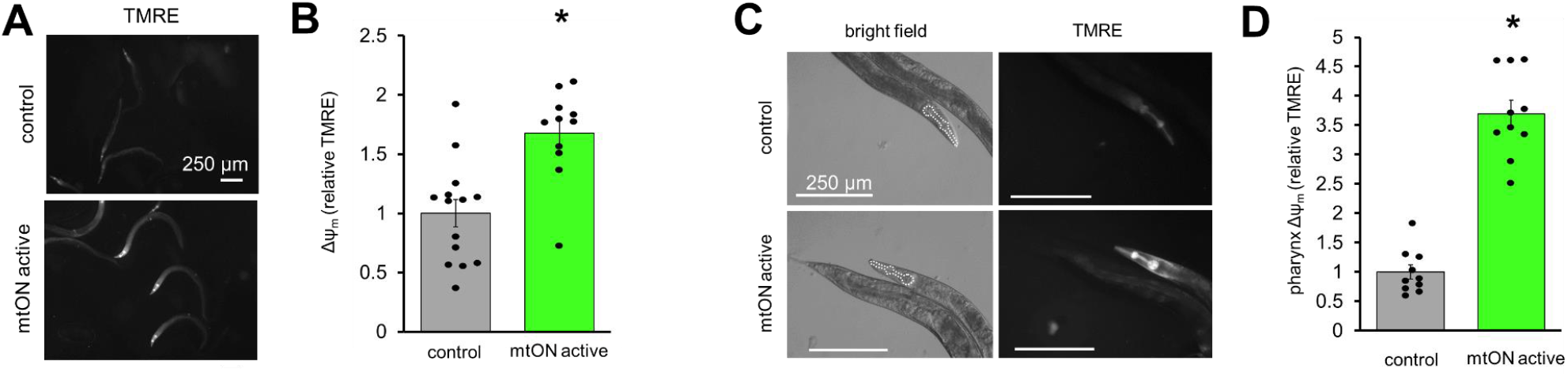
Single-copy expression of mtON increased Δψ_m_ in vivo. A) Images of day 4 adult worms expressing mtON in single-copy, stained with TMRE. Control is –ATR +light, mtON active is +ATR +light. Scale bar is 250 μm. B) Quantification of panel A, relative TMRE fluorescence of whole animals. Two tailed unpaired t test, *p < 0.05, n =14, 11 for control and mtON-active, respectively. Data are means + SEM, dots are individual worms. C) Images of day 4 adult animals expressing mtON in single-copy, stained with TMRE. Regions of interest (ROI) are drawn in the bright field panels around the pharynx, a mitochondria-rich tissue. Scale bar is 250 μm in all images. D) Quantification of panel C, relative TMRE fluorescence of the pharynx in adult animals. Two tailed unpaired t test, *p < 0.05, n = 10 animals for each condition. Data are means + SEM, dots are individual worms.

**Supplementary Figure 4.**
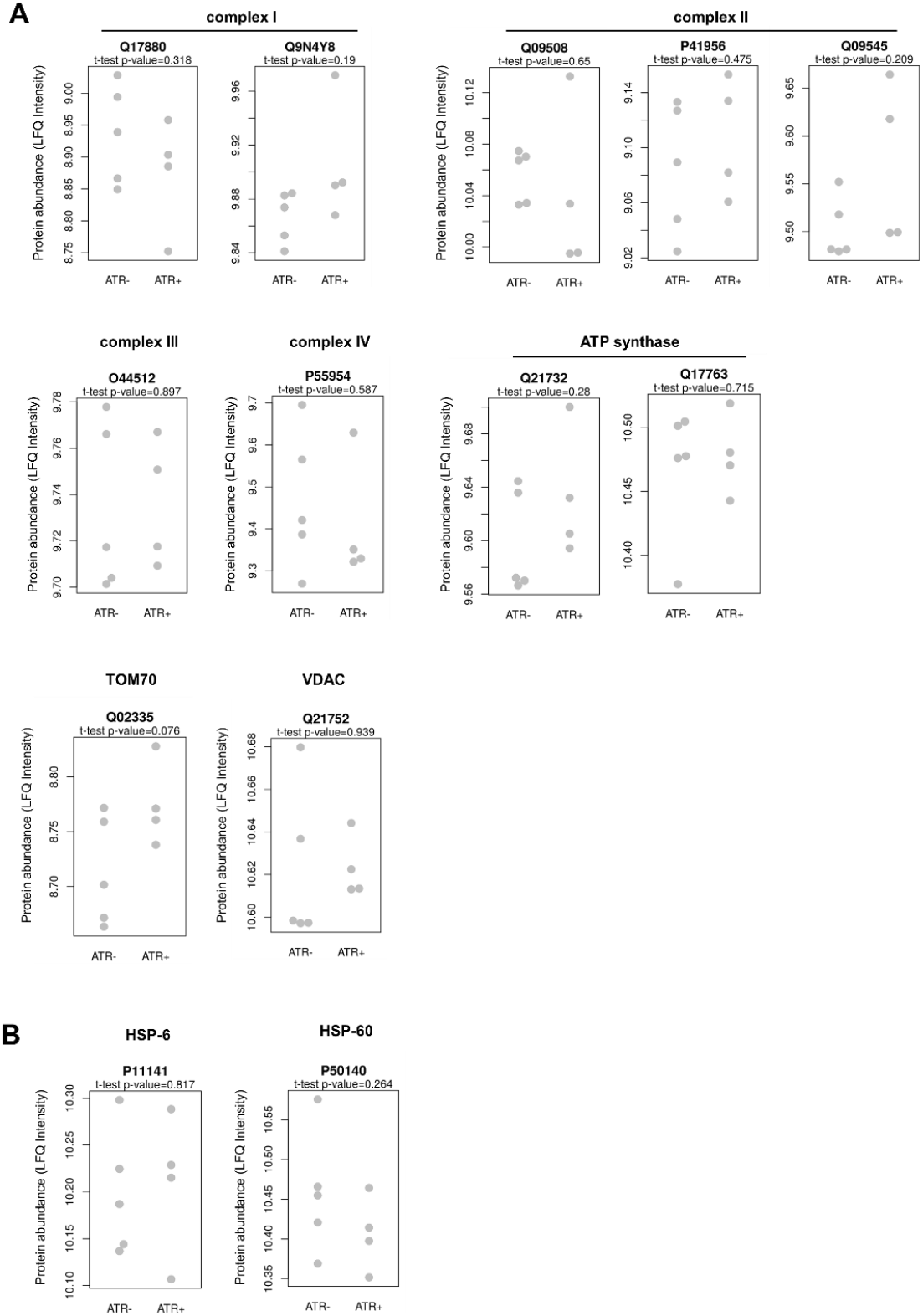
mtON activation did not affect mitochondrial mass or mitochondrial unfolded protein response by quantitative proteomics. A) The levels of key mitochondrial proteins were not significantly altered by 4 days of mtON activity, two-tailed unpaired t test, -ATR n = 5, +ATR n = 4 biological replicates. Genes encoding proteins are as follows. Complex I: *nuo-1, nuo-5.* complex II: *sdha-1, mev-1, sdhb-1.* Complex III: *isp-1.* Complex IV: *cox-5a.* ATP synthase: R04F11.2, *atp-5.* Translocase of the outer membrane (TOM70): *tomm-70.* VDAC: *vdac-1.* Uniprot *C. elegans* proteome database protein codes are presented above each plot. B) Proteins that mediate the mitochondrial unfolded protein response are not affected by mtON activation, two-tailed unpaired t test, -ATR n = 5, +ATR n = 4 biological replicates. Genes encoding proteins are *hsp-6* and *hsp-60.*

**Supplementary Figure 5.**
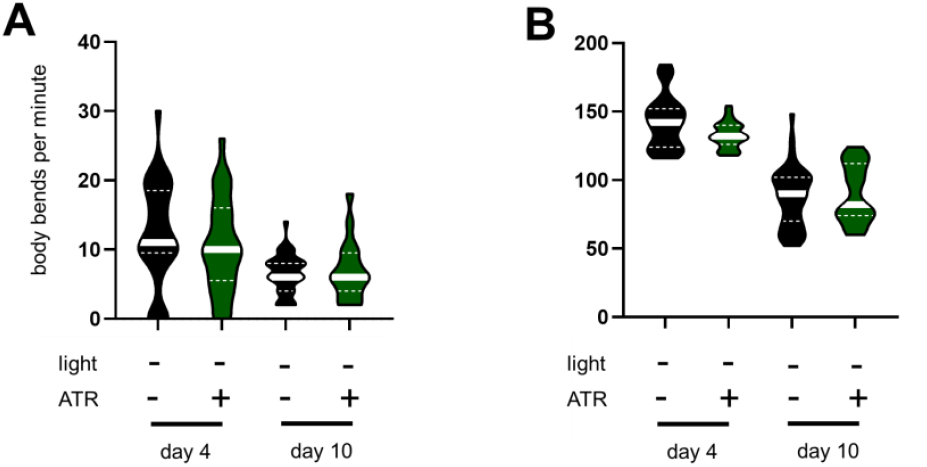
ATR treatment did not affect locomotion or thrashing with age. A) ATR treatment did not affect locomotion on solid media, and locomotion decreased with age. One-way ANOVA with Tukey’s test, n = 30, 30, 40, 40 animals for each violin from left to right. Data are medians + quartiles. B) ATR treatment did not affect thrashing in liquid, and thrashing decreased with age. One-way ANOVA with Tukey’s test, n = 31, 31, 43, 39 animals for each violin from left to right. Data are medians + quartiles. All statistical comparisons are presented in Supplementary Table 2.

**Supplementary Figure 6.**
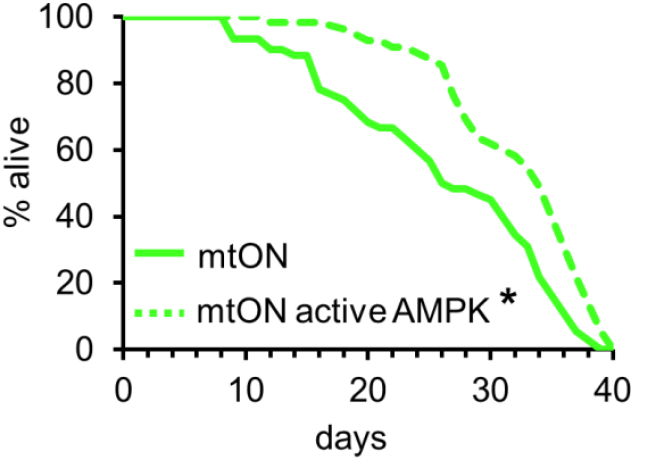
mtON increases lifespan of animals with constitutively active AMPK signaling. Survival curves of mtON-expressing animals versus mtON-expressing animals in a long-lived genetic background containing a constitutively active AMPK mutation [1]. Light treatment began at day 1 of adulthood. mtON activation in the CA-AMPK strain extended lifespan further compared to mtON alone, log-rank (Mantel-Cox) test, *p < 0.0001. Detailed statistical information is presented in Supplementary Table 1.

**Supplementary Table 1.** mtON reproducibly extended lifespan. We used two different strains of worms - APW32 expresses mtON as an extrachromosomal array (mtON ext. chr. array), and APW273 expresses mtON in single copy through CRISPR/Cas9 genome integration (mtON CRISPR). See methods section for genotype details. Light intensity was varied to maintain temperature. Bolded experiments correspond to the data displayed in Figure 2 and in Supplementary Figure 6.

| Experimental strain | Treatment | Light intensity (mW/cm<sup>2</sup>) | Median lifespan (days) | compared to: | p value treated vs. untreated (log-rank (Mantel-Cox) test) | % Lifespan extension | # of animals | FUDR |
| --- | --- | --- | --- | --- | --- | --- | --- | --- |
| mtON (ext. chr. array) | no treatment |  | 13 |  |  |  | 47 | No |
|  | ATR |  | 12 | no treatment | >0.05 |  | 46 | No |
|  | light | 2.10 | 13 |  |  |  | 47 | No |
|  | ATR, light | 2.10 | 17 | light | 0.035 | 31 | 43 | No |
| mtON (ext. chr. array) | no treatment |  | 14 |  |  |  | 59 | No |
|  | ATR |  | 14 | no treatment | >0.05 |  | 52 | No |
|  | light | 1.80 | 13 |  |  |  | 56 | No |
|  | ATR, light | 1.80 | 18 | light | 0.019 | 38 | 56 | No |
| mtON (CRISPR) | no treatment |  | 15 |  |  |  | 51 | No |
|  | ATR |  | 15 | no treatment | >0.05 |  | 51 | No |
|  | light | 0.15 | 15 |  |  |  | 46 | No |
|  | ATR, light | 0.15 | 16 | light | 0.026 | 6 | 54 | No |
| mtON (CRISPR) | no treatment |  | 15 |  |  |  | 52 | No |
|  | ATR |  | 17 | no treatment | 0.023 | 13 | 28 | No |
|  | light | 0.010 | 14 |  |  |  | 49 | No |
|  | ATR, light | 0.010 | 19 | light | <0.001 | 36 | 27 | No |
| mtON (CRISPR) | **no treatment** |  | **21** |  |  |  | **56** | **Yes** |
|  | **ATR** |  | **21** | **no treatment** | **> 0.05** |  | **53** | **Yes** |
|  | **light** | **0.15** | **17** |  |  |  | **53** | **Yes** |
|  | **ATR, light** | **0.15** | **22** | **light** | **< 0.0001** | **29** | **57** | **Yes** |
| N2 | No FCCP |  | **19** |  |  |  | 53 | Yes |
| | FCCP (10 μM) | | **19** | No FCCP | > 0.05 | | 36 | Yes |
| **mtON (CRISPR)** | **ATR, light, FCCP (10 μM)** | **0.15** | **21** |  |  |  | **61** | **Yes** |
|  | **ATR, light** | **0.15** | **28** | **ATR, light, FCCP** | **< 0.0001** | **33** | **43** | **Yes** |
| **mtON(CRISPR) vs. mtON(CRISPR) /CA-AMPK** | **mtON ATR, light** | **0.15** | **26** |  |  |  | **55** | **Yes** |
|  | **mtON/CA-AMPK ATR, light** | **0.15** | **34** | **mtON ATR, light** | **< 0.0001** | **31** | **55** | **Yes** |
| **mtON(CRISPR) /CA-AMPK** | Light | 0.15 | 25 |  |  |  | 63 | Yes |
|  | ATR, light | 0.15 | 28 | mtON/CA-AMPK Light | < 0.0001 | 12 | 63 | Yes |

**Supplementary table 2.** Statistical information. Only comparisons with p values less than 0.05 are shown. All tests are two sided.

| **Figure** | **test** | **Condition** | **n** | **comparison** | **p value** |
| --- | --- | --- | --- | --- | --- |
| 1C | One-way ANOVA with tukey's multiple comparisons test | day 1 | 44 | day 1 vs day 4 | < 0.0001 |
|  |  | day 4 | 18 | day 1 vs day 10 | 0.002 |
|  |  | day 10 | 30 |  |  |
| 1D | One-way ANOVA with tukey's multiple comparisons test | no ATR no light | 33 | no ATR light vs ATR light | 0.0003 |
|  |  | ATR no light | 34 |  |  |
|  |  | no ATR light | 26 |  |  |
|  |  | ATR light | 35 |  |  |
| 2C | One-way ANOVA with tukey's multiple comparisons test | Day 4 no ATR light | 30 | Day 4 no ATR light vs day 10 no ATR light | < 0.0001 |
|  |  | Day 4 ATR light | 30 | Day 4 no ATR light vs day 10 ATR light | < 0.0001 |
|  |  | Day 10 no ATR light | 40 | Day 4 ATR light vs day 10 no ATR light | < 0.0001 |
|  |  | Day 10 ATR light | 40 | Day 4 ATR light vs day 10 ATR light | < 0.0001 |
| 2D | One-way ANOVA with tukey's multiple comparisons test | Day 4 no ATR light | 32 | Day 4 no ATR light vs day 4 ATR light | < 0.0001 |
|  |  | Day 4 ATR light | 32 | Day 4 no ATR light vs day 10 no ATR light | < 0.0001 |
|  |  | Day 10 no ATR light | 40 | Day 4 no ATR light vs day 10 ATR light | < 0.0001 |
|  |  | Day 10 ATR light | 40 | Day 4 ATR light vs day 10 no ATR light | < 0.0001 |
|  |  |  |  | Day 4 ATR light vs day 10 ATR light | < 0.0001 |
|  |  |  |  | Day 10 no ATR light vs day 10 ATR light | 0.0020 |
| S2G | One-way ANOVA with tukey's multiple comparisons test | no ATR no light | 33 | no ATR no light vs ATR light | 0.0066 |
|  |  | ATR no light | 34 | ATR no light vs ATR light | 0.0061 |
|  |  | no ATR light | 26 | no ATR light vs ATR light | 0.0016 |
|  |  | ATR light | 35 |  |  |
| S3B | Two tailed unpaired t test | no ATR light | 14 | no ATR light vs ATR light | 0.0005 |
|  |  | ATR light | 11 |  |  |
| S3D | Two tailed unpaired t test | no ATR light | 10 | no ATR light vs ATR light | < 0.0001 |
|  |  | ATR light | 10 |  |  |
| S5A | One-way ANOVA with tukey's multiple comparisons test | Day 4 no ATR no light | 30 | Day 4 no ATR no light vs day 10 no ATR no light | 0.0001 |
|  |  | Day 4 ATR no light | 30 | Day 4 no ATR no light vs day 10 ATR no light | 0.0004 |
|  |  | Day 10 no ATR no light | 40 | Day 4 ATR no light vs day 10 no ATR no light | 0.0149 |
|  |  | Day 10 ATR no light | 40 | Day 4 ATR no light vs day 10 ATR no light | 0.0394 |
| S5B | One-way ANOVA with tukey's multiple comparisons test | Day 4 no ATR no light | 31 | Day 4 no ATR no light vs day 10 no ATR no light | < 0.0001 |
|  |  | Day 4 ATR no light | 31 | Day 4 no ATR no light vs day 10 ATR no light | < 0.0001 |
|  |  | Day 10 no ATR no light | 43 | Day 4 ATR no light vs day 10 no ATR no light | < 0.0001 |
|  |  | Day 10 ATR no light | 39 | Day 4 ATR no light vs day 10 ATR no light | < 0.0001 |

**Supplementary table 3.**
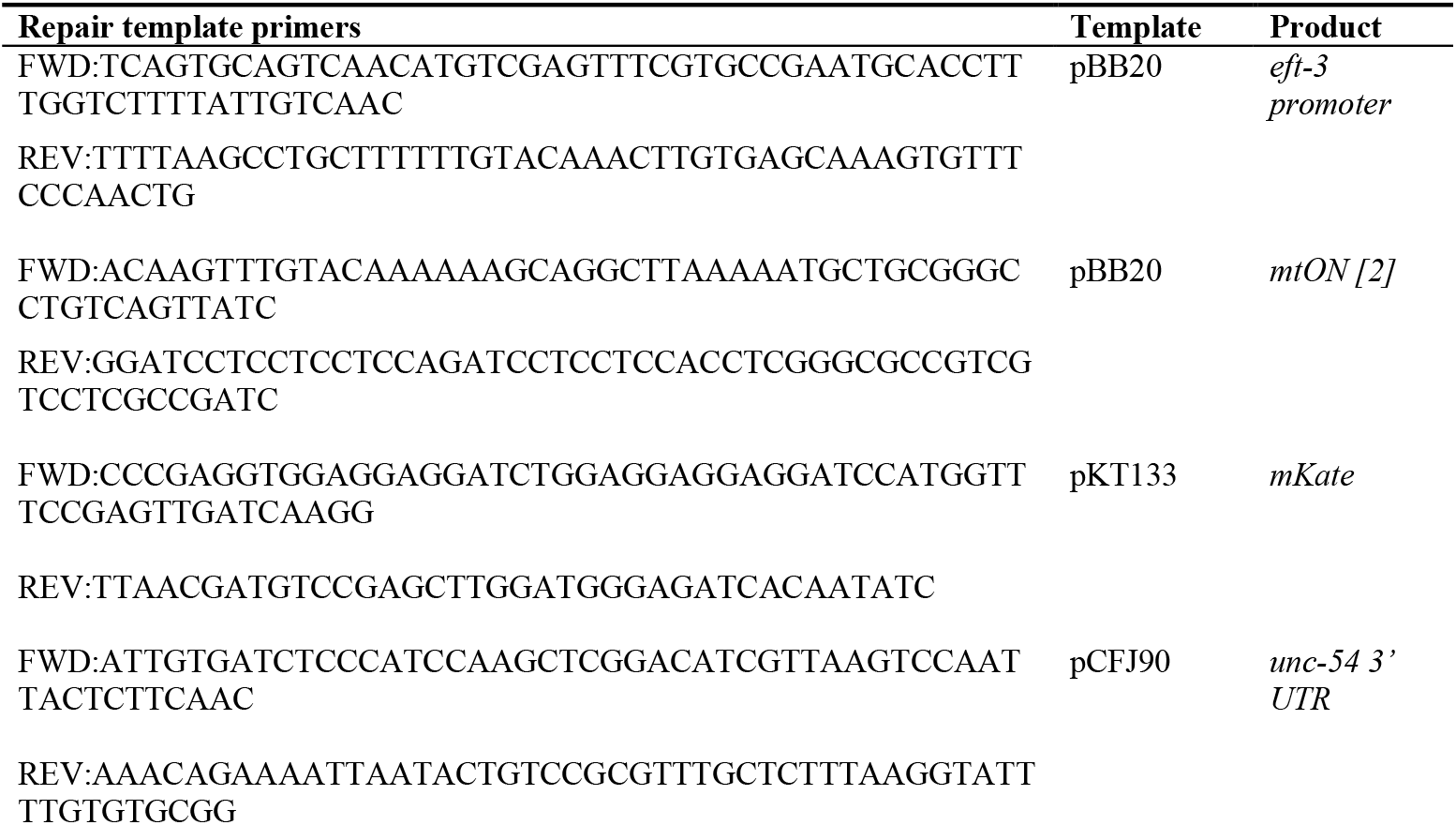
PCR repair template fragments

## Methods

### C. elegans *Strains and Maintenance*

Nematode growth medium (NGM) was used for *C. elegans* culture, and all maintenance and experiments were carried out at 20 °C. OP50 *E. coli* was used as a food source for all experiments. Where indicated, all-trans retinal (ATR) and/or FCCP was added to the food for a final concentration of 100 μM and 10 μM respectively, accounting for the volume of the NGM. Egg-lay synchronized day 1 adult hermaphrodite animals were used for all experiments unless otherwise noted. APW32 (genotype: *pha-1(e2123ts)* III; *jbmEx11 [pBJB20(Peft-3::Mitofilin(N’ 187aa)::Mac::GFP),* pC1 *(pha-1(+))])* expresses mtON as an extrachromosomal array [2]. APW273 (genotype: jbmSi10[eft-3p::Mitofilin(187N’aa)::Mac:mKate::unc-54 3’UTR *cxTi10816] IV) was created using Mos1 Element-Mediated CRISPR Integration approach [3]. Briefly, PCR fragments (Supplementary Table 3) were amplified and incorporated into a *Mos1* element on chromosome IV using CRISPR/Cas9 Homology-directed repair [4]. *C. elegans* were injected with a mix containing 25 mM KCl, 7.5 mM HEPES, 4 μg/μL tracrRNA, 0.8 μg/μL *Mos1* crRNA2 (target sequence: GTCCGCGTTTGCTCTTTATT), 0.8 μg/μL *dpy-10* crRNA, 50 ng/μL *dpy-10* ssODN, 2.5 μg/μL purified Cas9, and 300-400 pmol/μL of each PCR repair template fragment. APW312 (genotype: *jbmSi10* IV; *ulhIs248)* expresses a constitutively active AAK-2 with mtON and was generated by crossing WBM60 (genotype: *uthIs248 [Paak-2::aak-2 genomic(aa 1-321 with T181D)::GFP::unc-54 3’UTR, Pmyo-2::tdTomato]* with APW273.

### Whole Organism Mitochondrial Membrane Potential Measurement

Animals were stained for 24 hours with both 100 nM TMRE and 12 μM MitoTracker Green FM. TMRE was dissolved in ethanol and placed onto seeded plates, and MitoTracker Green FM was dissolved in DMSO and added to the OP50 food. Final concentrations accounted for the entire plate NGM volume. TMRE and MitoTracker Green FM were used to measure mitochondrial membrane potential and mitochondrial mass, respectively. Staining began at day 4 of adulthood, and animals were transferred to plates without dye for 1 hour prior to imaging to clear the gut of residual dye. Animals were mounted on 2% agarose pads under tetramisole (0.1% w/v) anesthesia. Texas Red and GFP filter sets were used to record images on an epifluorescence microscope (Nikon MVX10). Images were recorded with a Lumenera camera and associated software (Infinity Analyze). Fluorescence intensity was quantified using ImageJ by drawing regions of interest around individual animals, around the head region alone, or around individual pharynxes where indicated to determine their fluorescence intensity. Background signal was averaged and manually subtracted using ImageJ. Data are from 3 different experimental days. Confocal images were acquired using a Leica SP8X DMI6000 confocal microscope using a 63x oil immersion objective and a tunable white light laser (470-670 nn). Images were analyzed and prepared using Leica LASX Expert software and ImageJ.

### mtON Activation

Illumination was carried out with a 590 nm LED array with STOmk-II stimulator by Amuza placed 2 cm away from the surface of NGM plates. Intensity was measured using a calibrated optical power meter (1916-R, Newport Corporation). Animals were exposed to 1 Hz, 0.01 – 2.1 mW/mm^2^ light in order to maintain temperature across experiments. Note that under all lighting conditions mtON is maximally activated (lower limit of 0.01 mW/mm^2^ [2]). Lifespans were carried out starting at day 1 of adulthood until death or until they were removed to measure mitochondrial parameters. A digital temperature probe was used to adjust light intensity to maintain temperature at 20 °C under lighted conditions.

### Whole organism respiration

Oxygen consumption rate was measured using a Clark-type oxygen electrode (S1 electrode disc, DW2/2 electrode chamber, and Oxy-Lab control unit, Hansatech Instruments, Norfolk UK). Around 1000 animals per condition, per replicate were collected in M9 allowed to settle by gravity, rinsed in M9 buffer, settled again, and finally added to the electrode chamber in 0.5 mL of continuously stirred M9 buffer. FCCP was added at 160 μM final concentration in the chamber to induce maximal respiration. Respiration rates were measured for 10 minutes or until stable. Animals were then collected in M9 buffer for protein quantification using the Folin-phenol method.

### Lifespan Analysis

Animals were transferred to new plates every 2 days until reproduction ceased, and as necessary to replenish food. Where indicated, 50 μM FUDR was used in the NGM to prevent progeny from developing. Animals that did not move in response to a light touch to the head with a platinum wire were scored as dead and removed from assay plates. 1 - 3 plates for each condition were scored concurrently with ~15-70 animals per plate. All experiments were performed to maintain 20° C which sometimes required adjustment of light intensity (noted in Supplementary Table 1). Animals were illuminated only during adulthood. FCCP was added to plates for 10 μM final concentration. This dose did not affect lifespan on its own (see supplementary table 1, N2 FCCP lifespan. All lifespans comprise 3 biological replicates pooled for each experiment.

### Locomotion Assays

Synchronized animals were observed and locomotion was scored by counting body bends according to previous methods [5]. Locomotion was scored in the presence of food on solid media. Thrashing was similarly analyzed with animals placed in M9 buffer to move freely in liquid. Body bends were counted for 30 seconds for each animal in all cases, and multiplied by 2 to represent body bends per minute in accordance with previously used protocols [2, 6]. Data are from at least 3 different experimental days.

### *Protein extraction from* C. elegans

*C. elegans* were washed and stored in water after 4 days of illumination. Worms were centrifuged for 5 minutes at 200x g at 4°C, supernatant was discarded. The worm pellet was then resuspended in 100μl Lyse (iST Sample Preparation Kit, Preomics, Planegg/Martinsried), incubated at 95°C for 10 minutes with 500 rpm shaking and sonicated 10x 10 seconds at 30% amplitude. The sample was then centrifuged at 8000x g for 15 minutes at 4°C and the supernatant was transferred to a new vial. The protein concentration was measured with the NanoDrop 2000.

### Proteomics sample preparation

100 μg protein in 50μl Lyse was recommended as starting material for the sample preparation with the Preomics iST Sample Preparation Kit. In the case the protein concentration was higher than 2μg/μl, the sample was diluted with Lyse. The preparation was performed according to the supplier guidelines. (Kit: iST Sample Preparation Kit, Preomics, Order nr. P.O.00001).

### LC-MS/MS data acquisition

LC-MS/MS measurements were performed on an Ultimate 3000 RSLCnano system coupled to a Q-Exactive HF-X mass spectrometer (Thermo Fisher Scientific). Peptides were delivered to a trap column (ReproSil-pur C18-AQ, 5 μm, Dr. Maisch, 20 mm × 75 μm, self-packed) at a flow rate of 5 μL/min in 100% solvent A (0.1% formic acid in HPLC grade water). After 10 minutes of loading, peptides were transferred to an analytical column (ReproSil Gold C18-AQ, 3 μm, Dr. Maisch, 450 mm × 75 μm, self-packed) and separated using a 110 min gradient from 4% to 32% of solvent B (0.1% formic acid in acetonitrile and 5% (v/v) DMSO) at 300 nL/min flow rate. The Q-Exactive HF-X mass spectrometer was operated in data dependent acquisition (DDA) and positive ionization mode. MS1 spectra (360-1300 m/z) were recorded at a resolution of 60,000 using an automatic gain control (AGC) target value of 3e6 and maximum injection time (maxIT) of 45 msec. Up to 18 peptide precursors were selected for fragmentation in case of the full proteome analyses. Only precursors with charge state 2 to 6 were selected and dynamic exclusion of 30 sec was enabled. Peptide fragmentation was performed using higher energy collision induced dissociation (HCD) and a normalized collision energy (NCE) of 26%. The precursor isolation window width was set to 1.3 m/z. MS2 Resolution was 15.000 with an automatic gain control (AGC) target value of 1e5 and maximum injection time (maxIT) of 25 msec.

### LC-MS/MS data analysis

Peptide identification and quantification was performed using MaxQuant (version 1.6.3.4). MS2 spectra were searched against the Uniprot *C. elegans* proteome database (UP000001940, 26672 protein entries, downloaded 21.12.2020) supplemented with the mKate-tagged proton pump protein plus common contaminants. Trypsin/P was specified as proteolytic enzyme. Precursor tolerance was set to 4.5 ppm, and fragment ion tolerance to 20 ppm. Results were adjusted to 1 % false discovery rate (FDR) on peptide spectrum match (PSM) level and protein level employing a target-decoy approach using reversed protein sequences. The minimal peptide length was defined as 7 amino acids, the “match-between-run” function was disabled. Carbamidomethylated cysteine was set as fixed modification and oxidation of methionine and N-terminal protein acetylation as variable modifications. The label free quantification (LFQ) [7] from MaxQuant was used to represent the relative abundance of proteins across samples. The ATP synthase, HSP6, HSP 60, TOM70, VDAC, mitochondrial complex proteins were manually selected. The different expression of these protein between the ATR positive and negative samples were performed using student’s t test.

## Data availability

All data are presented in the manuscript. Any raw data files will be made available upon request.

The proteomic raw data and MaxQuant search files will be deposited to the ProteomeXchange Consortium (http://proteomecentral.proteomexchange.org) via the PRIDE partner repository.

